# Gamma oscillations in primate primary visual cortex are severely attenuated by small stimulus discontinuities

**DOI:** 10.1101/2020.07.19.210922

**Authors:** Vinay Shirhatti, Poojya Ravishankar, Supratim Ray

**Affiliations:** Centre for Neuroscience, Indian Institute of Science, Bangalore, India, 560012; IISc Mathematics Initiative, Indian Institute of Science, Bangalore, India, 560012; Institute for Systems Research, University of Maryland, College Park, Maryland 20742; Committee on Computational Neuroscience, University of Chicago, Illinois 60637, Facsimile

**Keywords:** Gamma, LFP, V1, Gratings, Discontinuities

## Abstract

Gamma oscillations have been hypothesized to play an important role in feature binding, based on the observation that continuous long bars induce stronger gamma in the visual cortex than bars with a small gap. Recently, many studies have shown that natural images, that have discontinuities in several low-level features, do not induce strong gamma oscillations, questioning their role in feature binding. However, the effect of different discontinuities on gamma has not been well studied. To address this, we recorded spikes and local field potential from two monkeys while they were shown gratings with discontinuities in space, orientation, phase or contrast. Gamma, but not spiking activity, drastically reduced with small discontinuities in all cases, suggesting that gamma could be a resonant phenomenon. An excitatory-inhibitory population model with stimulus-tuned recurrent inputs showed such resonant properties. Therefore, gamma could be a signature of excitation-inhibition balance, which gets disrupted due to discontinuities.

## Introduction

Gamma oscillations (∼30-80 Hz) are strongly induced in the primary visual cortex (area V1) by stimuli such as gratings, bars or colors (Bartoli et al., 2020). One influential hypothesis posits that gamma oscillations play a role in visual perceptual grouping or feature binding, based on the finding that continuous bars induce stronger gamma synchronization between neurons whose receptive fields (RFs) contain parts of the bar as compared to discontinuous bars, even when the discontinuity is outside their RFs (Castelo-Branco et al., 1998; Engel et al., 1991; Gray et al., 1989; Gray and Prisco, 1997). However, in the case of natural images, studies have reported disparate observations regarding the consistency of gamma oscillations (Bartoli et al., 2019; Brunet et al., 2013; Hermes et al., 2014), casting doubts on a causal role for them in feature binding in a natural setting. Because such natural stimuli might occur as discontinuities along many feature dimensions across the RF of neurons, it is important to study how different types of structural irregularities affect firing responses and gamma oscillations. A recent study explored discontinuities in chromatic content and showed that stimulus discontinuities can reduce gamma synchronization between responsive neurons (Peter et al., 2019). However, the effect of discontinuities along other feature dimensions, such as orientation, phase and contrast, on gamma oscillations and firing rates remain largely unknown.

In addition, it is unclear how gamma oscillations depend on the size of the discontinuity. Recently, Hermes and colleagues proposed an image-computable model of gamma oscillations, in which the amplitude of gamma depends on the variability across orientation channels (Hermes et al., 2019). In such models, reduction in gamma is expected to be graded and proportional to the size of the discontinuity. However, other studies have suggested that gamma could be a resonant phenomenon arising due to a tight interplay of excitatory and inhibitory (E-I) signals in a neuronal network (Atallah and Scanziani, 2009; Brunel and Wang, 2003; Buzsáki and Wang, 2012; Jia et al., 2013; Moca et al., 2014; Veit et al., 2017) Indeed, the V1 RF structure has an excitatory center region flanked by suppressive near-surround and far-surround regions, and involves interactions between feedforward geniculocortical signals, lateral intracortical signals from horizontal connections and feedback signals from higher areas (Angelucci et al., 2017). Stimulus discontinuities could potentially modulate the interactions between these diverse neuronal sub-populations and alter the levels of E-I in this network, which may result in a drastic reduction in gamma even with a small discontinuity.

To address these questions, we recorded spikes and local field potential (LFP) from area V1 of passively fixating alert monkeys using microelectrode arrays, while they were shown sinusoidal luminance gratings with or without discontinuities that varied along one of four dimensions: space, orientation, phase and contrast. Further, the magnitude of discontinuity for each dimension was parametrically varied. We compared how gamma oscillations and firing rates changed with the magnitude of such discontinuities. Finally, we built an E-I network based on Wilson-Cowan model operating in an inhibition stabilized mode (Jadi and Sejnowski, 2014; Tsodyks et al., 1997; Wilson and Cowan, 1972) and added stimulus dependent local recurrent inputs to model discontinuities. This simple model could mimic crucial aspect of our observations.

## Results

We implanted a microelectrode array in area V1 of two monkeys and estimated the RFs of the recorded sites by flashing small sinusoidal luminance gratings on locations forming a dense rectangular grid on the approximate aggregate RF for all the sites ((Dubey and Ray, 2019, 2016); see Methods for details). We have previously shown that large gratings induce two distinct gamma oscillations in V1, termed slow (20-35 Hz) and fast (35-70 Hz) gamma (Murty et al., 2018). Here, we presented large static gratings (radii of 9.6° and 6.4° for the two monkeys) at a spatial frequency of 4 cycles/degree (cpd), 100% contrast (except in the contrast discontinuities experiment), at an orientation that induced strong fast gamma oscillations (although for Monkey 2, these induced moderately strong slow gamma as well), and introduced discontinuities of different types. Therefore, the following results are focused on the fast gamma band, and ‘gamma’ refers to this band. Unless otherwise stated, for the discontinuous gratings, the radius of the inner grating was fixed at 0.3° and 0.2° for Monkeys 1 and 2 respectively, which was close to the average RF sizes (mean ± s.e.m. for Monkey 1: 0.28° ± 0.009°, Monkey 2: 0.17° ± 0.0065°). Thus, the discontinuity across experiments occurred approximately in the visual space corresponding to a transition between the center and surround. In each session, stimuli were centered on the RF center of one of the recorded sites.

### Annular cut discontinuity disrupts gamma

We first tested the effect of a discontinuity in space by introducing an annular cut in the gratings, whose width could take one of the following 5 values: 0° (no discontinuity), 0.025°, 0.05°, 0.1°, and 0.2° (Figure 1A, topmost row). The trial averaged time frequency (TF) difference spectra for an example site from a session in Monkey 1 shows a drastic reduction in gamma power by ∼58% with an introduction of the smallest tested discontinuity of 0.025° (Figure 1A, second row). This effect was qualitatively consistent in the population across multiple sessions in both monkeys (Figure 1A, rows 3 and 4) and was also evident in the average change in power plot (computed between a stimulus period of 250-750 ms and compared against a baseline period of −500 to 0 ms, where 0 ms represents stimulus onset) for different stimuli (Figure 1B). Here, we also observed a second peak at ∼80-100 Hz in Monkey 1, which was simply a harmonic of the fast gamma. In comparison, the spiking activity showed only a modest increase across different conditions (Figure 1C) as the surround suppression reduced with increasing annular cut width.

**Figure 1:**
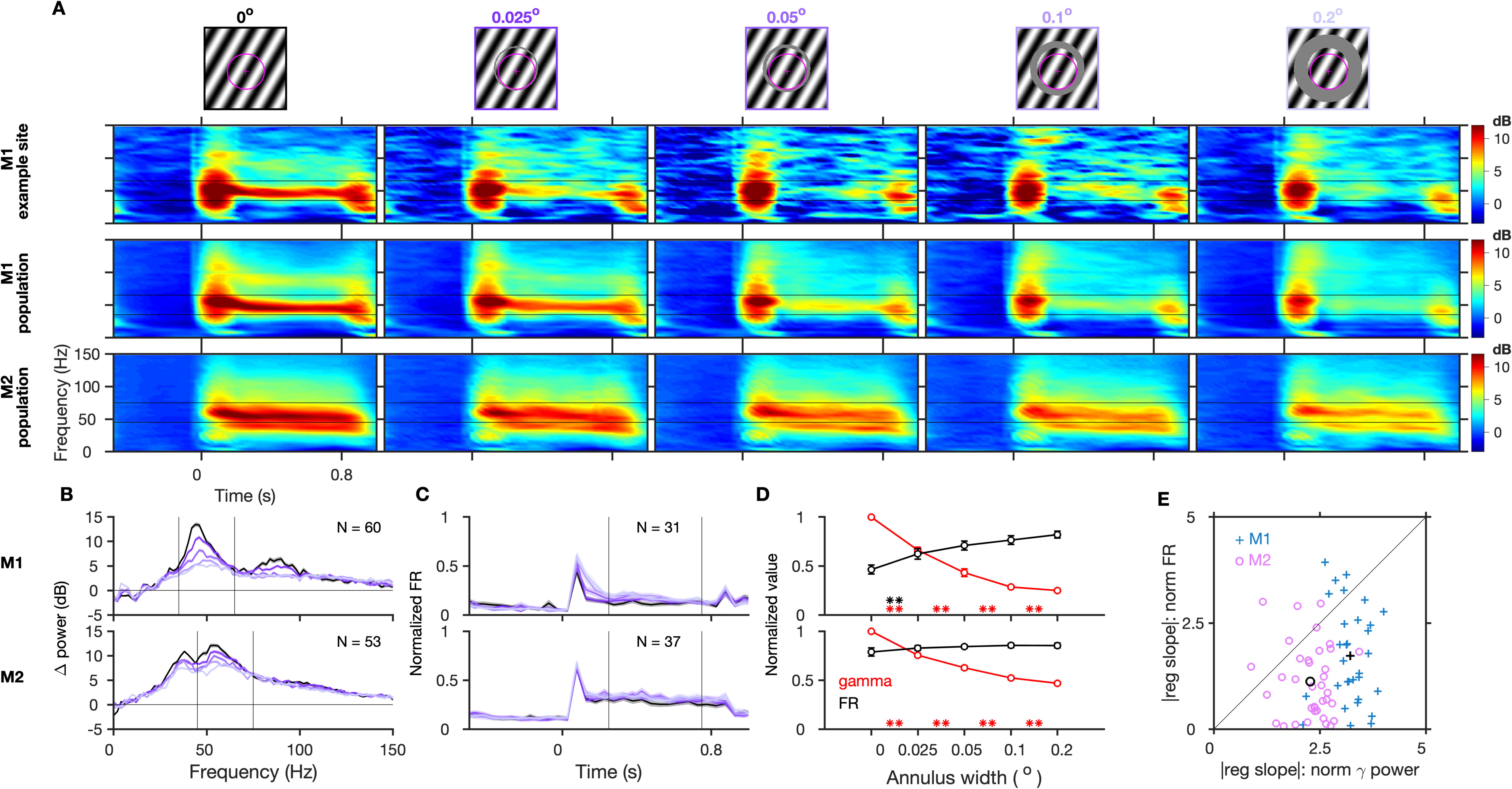
Gamma oscillations are reduced by annular discontinuity. A) Top row: grating stimuli with annular cut discontinuity along with the receptive field (magenta) of an example center site in Monkey 1 from an example session. Corresponding cut width is mentioned and color coded for other plots. Trial-averaged TF difference spectra for the example center site in Monkey 1 (2^nd^ row), and for the population (averaged across center electrodes and sessions) in Monkey 1 and 2 (following rows; number of sites, N = 60 and 53 respectively). Gratings were presented between 0 to 0.8 seconds. B) The corresponding mean change in power from baseline ([-0.5 0] s) to stimulus period ([0.25 0.75] s), averaged across these electrodes and sessions. Black lines in A) and B) indicate the gamma ranges (35-65 Hz and 45-75 Hz for the two monkeys). These ranges are chosen to avoid the slow gamma, which is prominent in Monkey 2. C) Mean normalized firing rate (averaged across selected spiking center electrodes and sessions) for different conditions. The firing rate for each electrode is normalized by the maximum for that electrode across time and conditions. D) Normalized gamma power and firing rate during the stimulus period (black lines in (C); normalized for each selected spiking center electrode in (C) by the maximum across conditions, averaged across these center electrodes across sessions) for different annular discontinuity values. The error bars denote the standard error of mean (s.e.m.); and the asterisks in D) indicate statistical significance (** for p<0.01, * for p<0.05, WSR test) of change in normalized gamma (red) or firing rate (black) between the flanking values of annulus widths. E) Magnitude of slope of regression with annulus width (Δvalue/degree of visual angle) of normalized gamma power versus corresponding slopes for normalized firing rate. Each data point represents a selected center electrode. Black data points indicate the mean value across these electrodes.

To directly compare the relative effects of annular discontinuity on gamma power and spiking activity at the population level, we first selected units which had a mean firing rate of at least 1 spike/sec during the stimulus epoch for at least one of the stimulus conditions, normalized both gamma power and firing rates by dividing by the maximum value across the five stimulus conditions, and computed the mean across sites (Figure 1D). While the mean normalized gamma power decreased significantly for each discontinuity level (as indicated by red asterisks in Figure 1D) in both monkeys, the increase in firing rate was less salient, only reaching significance for the comparison between no discontinuity and the smallest discontinuity in Monkey 1. Slopes of regression of change in mean normalized gamma power with increasing annulus width were −3.22 ± 0.08 units/° (mean ± s.e.m. units per degree of visual angle) and −2.27 ± 0.09 units/° for the two monkeys, both significantly negative (Monkey 1: p = 0.12 × 10^−5^, Wilcoxon signed rank (WSR) test, Monkey 2: p = 0.11 × 10^−6^), whereas similar regression slopes with normalized firing rates were 1.52 ± 0.25 units/° for Monkey 1 (p = 0.3 × 10^−4^, WSR test) and 0.26 ± 0.23 units/° for Monkey 2 (p = 0.41). Importantly, the magnitude of slope was significantly greater for gamma power than for firing rates across individual sites (Figure 1E, p = 0.12 × 10^−4^ and 0.77 × 10^−5^ for Monkey 1 and 2 respectively, WSR test), implying that the mean rate of change in normalized gamma with annulus width was significantly greater than that for normalized firing rates. These results indicate that overall gamma was more sensitive to the annular discontinuity than firing rate.

To test whether our results depended on the location of the discontinuity, we performed the same experiment but with the discontinuity appearing at one of the following 4 radii from the center of the stimulus: 0.15°, 0.3°, 0.6° and 1.2° in Monkey 1 and 0.1°, 0.2°, 0.4° and 0.8° in Monkey 2. For each case, the width of the cut could take one of the same 5 values as in the annular cut experiment. Figure 2A and 2B show the normalized gamma power and firing rates for the two monkeys (see Supplementary Figure 1 for more details). Gamma was more severely disrupted by a discontinuity that occurred closer to the center (Figure 2A, Supplementary Figure 1A-1B). This was reflected in the magnitude of the regression slope between normalized gamma power and annulus width, which decreased for farther cut locations (Figure 2C). On the other hand, increases in firing rate were more modest (Figure 2B, Supplementary Figure 1C), with regression slopes significantly smaller than the corresponding slopes for gamma for smaller cut radii (as indicated in Figure 2C).

**Figure 2:**
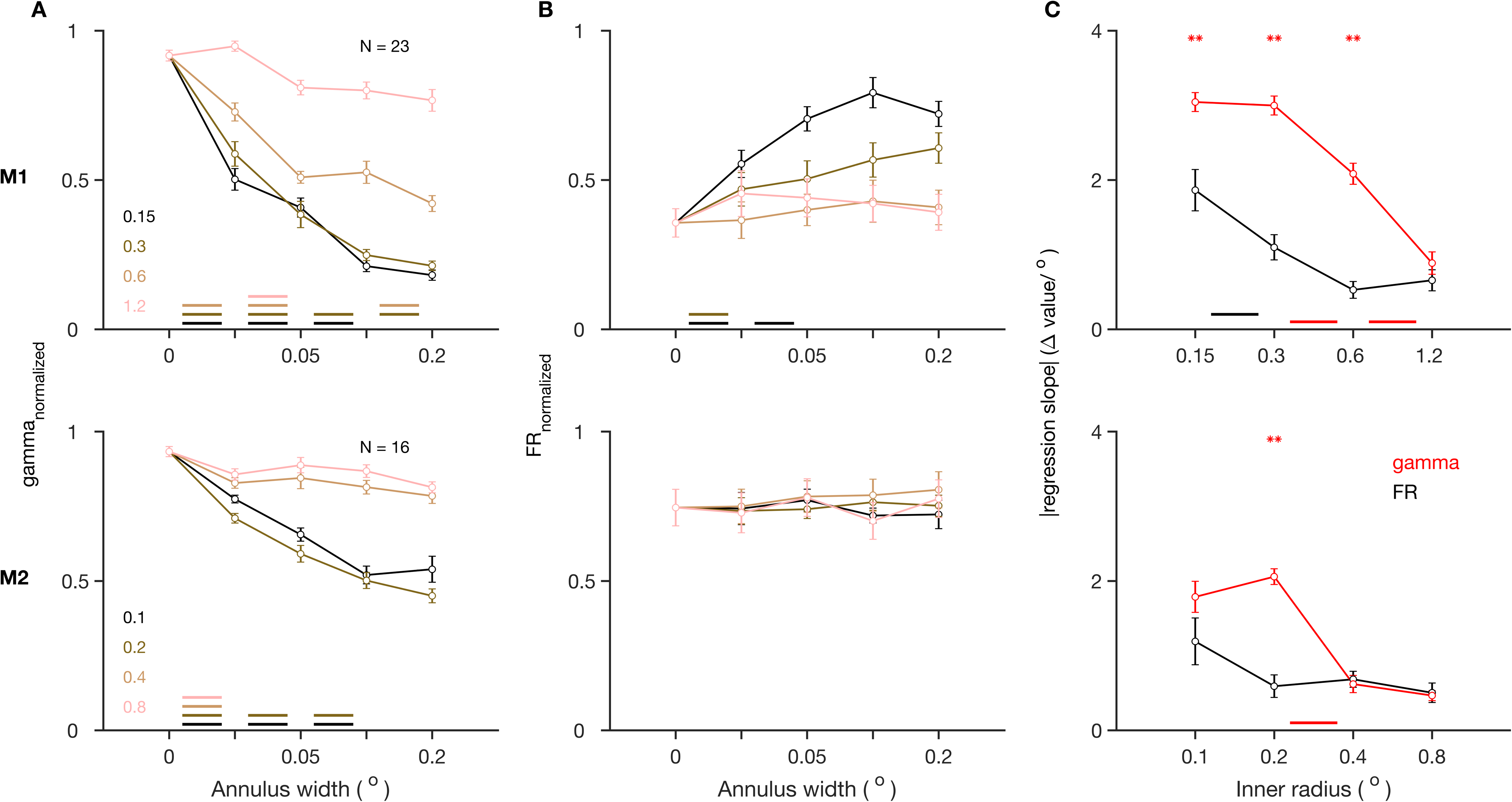
Effect of annular discontinuity at different locations. A) Normalized gamma power and B) Normalized firing rate, for stimuli with annular discontinuity at different inner radii from the center of the stimulus, and as a function of the width of annular discontinuity, averaged across center electrodes, for Monkey 1 (top) and 2 (bottom). C) Magnitude of slope of regression of the values in A and B on annulus width for different inner radii, averaged across the center electrodes. The color-coded lines at the bottom indicate statistical significance (p<0.01, WSR test) of change in corresponding value between the flanking values of annulus widths. The asterisks at the top indicate statistical significance (p<0.01, WSR test) of these values being greater for normalized gamma than for normalized firing rates.

### Effect of orientation discontinuity

To evaluate the effect of orientation discontinuity we varied the relative orientation of the inner and outer parts of the static grating stimulus (Figure 3A). Average TF difference spectra across trials of the different stimuli, for an example site from a session in Monkey 1, show that gamma oscillations were strongest for matched orientations, and their strength reduced drastically even with the smallest mismatch of 10° on both sides (by ∼72% for a difference of −10° between outer and inner orientation ((*O-I*)°), and by ∼69% for (*O-I*)° = 10°). As the orientation difference systematically increased between the inner and outer gratings, there was a drastic reduction in the strength of gamma power across the population in both monkeys as evidenced in the average TF spectra (Figure 3A, rows 3 and 4) and change in power (Figure 3B). On the other hand, spiking activity increased as the mismatch between inner and outer orientation increased (normalized firing rates, Figure 3C). This is consistent with previous studies of cross orientation suppression where lateral inhibition from the surround has been shown to be the strongest when the center and surround orientation are matched and weakens with increase in orientation differences (Blakemore and Tobin, 1972; Knierim and van Essen, 1992; Sillito et al., 1995).

**Figure 3:**
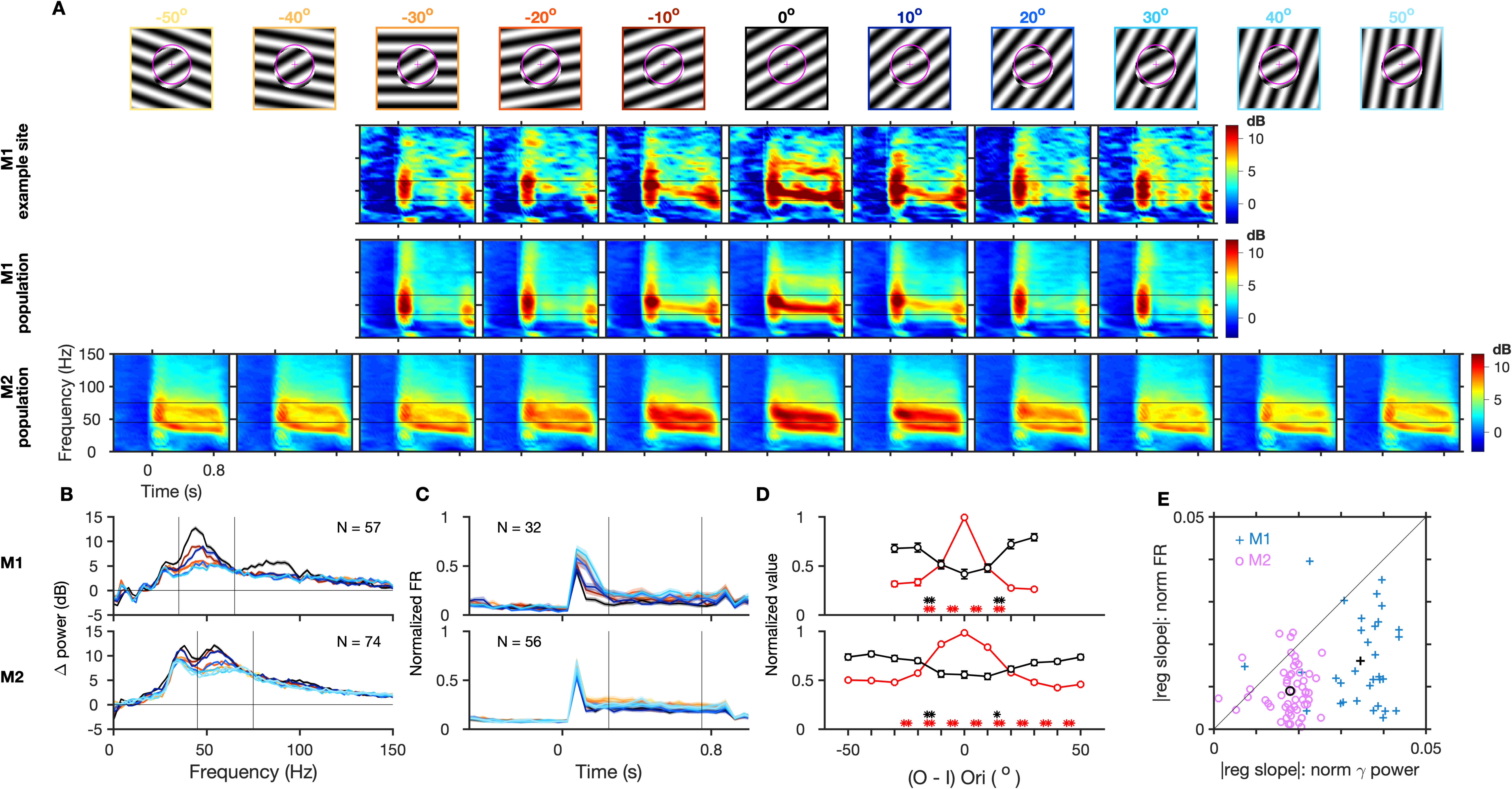
Gamma oscillations are reduced by orientation discontinuity. Same format as in Figure 1. For Monkey 2 there are two additional steps of discontinuity on either side (−50° to 50° in steps of 10°).

For a subset of strongly firing units, we compared the dependence on orientation discontinuity of the normalized gamma power to normalized spiking activity (Figure 3D), similar to the analysis in Figure 1D. For the smallest orientation discontinuity that we presented, normalized gamma power reduced significantly for both sides by ∼48% (for (*O-I)*° = −10°, p = 0.96 × 10^−6^, WSR test) and ∼52% (for (*O-I)*° = 10°, p = 0.80 × 10^−6^) in Monkey 1 and ∼12% (for (*O-I)*° = −10°, p = 0.67 × 10^−7^) and ∼15% (for (*O-I)*° = 10°, p = 0.13 × 10^−8^) in Monkey 2, whereas firing rates remained mostly unchanged (Figure 3D, WSR test, p = 0.06 and p = 0.29 for the two sides in Monkey 1; similarly p = 0.64 and p = 0.43 in Monkey 2). To further compare the rate of change in gamma and firing rates due to orientation discontinuity, we computed the slope of regression of their normalized values with discontinuity on both sides over the range where mean values varied clearly ((*O-I)*° = 0° to ±20° in Monkey 1, and (*O-I)*° = 0° to ±30° in Monkey 2). Across sites the slopes were significantly negative for gamma (p = 0.88 × 10^−6^ and 0.80 × 10^−6^ on the two sides in Monkey 1, and 0.75 × 10^−10^ and 0.89 × 10^−10^ in Monkey 2, WSR test), and significantly positive for firing rates (p = 0.16 × 10^−4^ and 0.54 × 10^−4^ in Monkey 1, p = 0.64 × 10^−4^ and 0.48 × 10^−2^ in Monkey 2, WSR test). However, the magnitude of the mean slope (averaged across both sides) was larger for gamma than for firing rates consistently across individual sites in both the monkeys (Figure 3E, p = 0.42 × 10^−5^ in Monkey 1, and 0.66 × 10^−8^ in Monkey 2, WSR test). Interestingly, the mean firing responses changed significantly between conditions of 10° to 20° orientation differences between the inner and outer gratings, after which they again remained comparable for increasing orientation differences in both the monkeys (Figure 3D). Gamma, on the other hand, was sensitive to orientation differences across a larger range and this effect was more consistent across the recorded sites.

### Effect of phase discontinuity

Next, we introduced increasing levels of spatial phase difference between the inner and outer gratings. Gamma dropped drastically with the smallest phase difference of 60° on both sides (both (*O-I)ϕ*° = 60° and 300° imply a phase difference of 60° from two sides of the spatial cycle) as seen in the average TF difference spectra for an example site in Monkey 1 (Figure 4A, row 2, gamma reduced by ∼57% for (*O-I)ϕ*° = 60° and ∼52% for (*O-I)ϕ*° = 300°) and for the population in both the monkeys (Figure 4A, 3^rd^ and 4^th^ row). Across the population, the matched condition showed the strongest gamma oscillations and the strength of gamma reduced progressively as the phase disparity increased from both sides of the spatial cycle until 180°, at which the center and surround were in perfect anti-phase, showing that gamma changed in accordance with the magnitude of discontinuity (Figure 4A, 4B). Firing rates remained mostly unchanged with the smallest discontinuities and showed a small increase with bigger discontinuities (Figure 4C, 4D).

**Figure 4:**
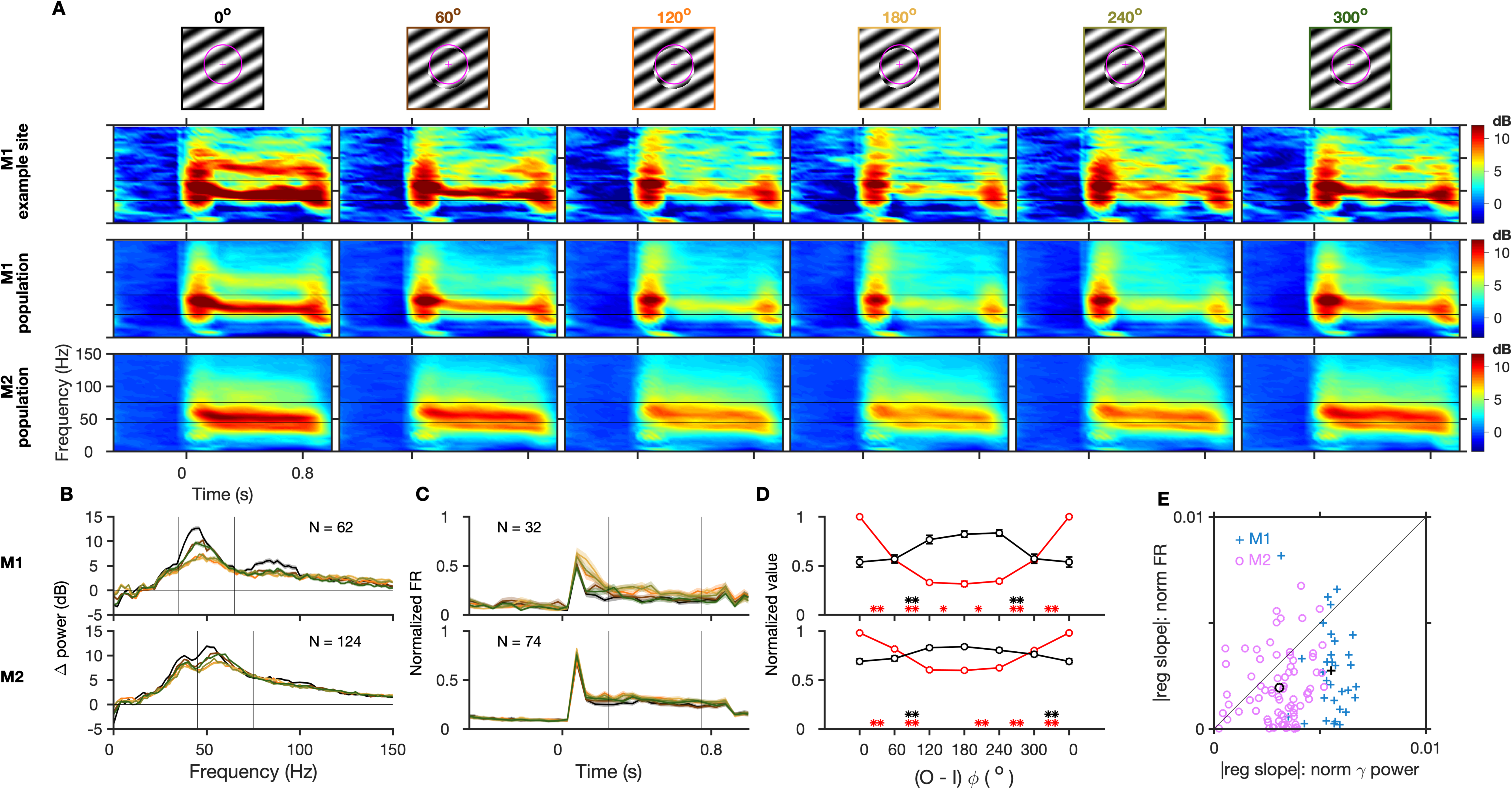
Gamma oscillations are reduced by phase discontinuity. Same format as in Figure 1 and 3. In (D) the values for 0° are repeated on the right side due to the circular scale of phases and for easy comparison with values for 300°, which is one of the two smallest discontinuities in this stimulus set.

The stimulus period mean normalized gamma power (Figure 4D, similar to Figure 1D and 3D) reduced significantly by ∼44% with the smallest phase differences on both sides in Monkey 1 (p = 0.40 × 10^−17^ for (*O-I)ϕ*° = 60° and 0.18 × 10^−14^ for (*O-I)ϕ*° = 300°, t-test) and by ∼17% ((*O-I)ϕ*° = 60°, p = 0.23 × 10^−13^) and ∼18% ((*O-I)ϕ*° = 300°, p = 0.61 × 10^−17^) in Monkey 2. The corresponding change in normalized firing rates was not significant in Monkey 1 (p = 0.36 and 0.48, t-test) and increased significantly only for (*O-I)ϕ*° = 300° in Monkey 2 (p = 0.57 × 10^−3^; p = 0.26 for (*O-I)ϕ*° = 60°). To compare the dependence of gamma and firing rates on phase discontinuity at each site, we computed the slope of regression of their normalized values with phase discontinuities from 0° to 180° from both sides of the spatial cycle. Slopes were significantly negative for normalized gamma (p = 0.8 × 10^−6^ for both sides in Monkey 1; p = 0.87 × 10^−13^ and 0.20 × 10^−12^ for Monkey 2, WSR test) and significantly positive for firing rates (p = 0.58 × 10^−4^ and 0.10 × 10^−2^ in Monkey 1; p = 0.62 × 10^−3^ and 0.12 × 10^−3^ in Monkey 2) on both sides in both monkeys. The magnitude of slope was significantly larger for gamma than for firing rates across individual sites (Figure 4E, p = 0.20 × 10^−4^ in Monkey 1; p = 0.33 × 10^−5^ in Monkey 2). Thus, change in gamma power was more sensitive to phase discontinuities for a much broader range compared to firing rate (Figure 4D) and showed a higher sensitivity consistently across sites (Figure 4E). The most significant changes in firing responses were observed for bigger mismatches in phases of 120° or 240° between the center and surround region. They did not increase further for larger mismatches in both monkeys (Figure 4D, black line). Thus, there is a critical sub-range of maximum sensitivity of firing rates to phase differences, similar to the orientation discontinuity effects.

### Effect of contrast discontinuity

Increasing the contrast of a visual stimulus has been shown to increase the spiking responses as well as the strength (Henrie and Shapley, 2005) and frequency of gamma oscillations in V1 (Jia et al., 2013; Ray and Maunsell, 2010). Likewise, increasing the stimulus size, which is expected to increase surround suppression, also increases gamma power but reduces the peak frequency (Gieselmann and Thiele, 2008; Jia et al., 2013; Ray and Maunsell, 2011). We studied how a discontinuity in the contrast profile across the inner and outer regions of the grating affects gamma oscillations, by varying their contrast independently over five different values (25 combinations). In Figure 5A, the inner and outer contrasts were matched along the diagonal, and contrast discontinuity progressively increased away from this diagonal. If gamma depended mainly on the overall input strength, it should increase with an increase in contrast of either the inner or outer segment of the stimulus. In such a scenario gamma should increase from the left to the right column (Figure 5A, as outer contrast increases) and from the top to the bottom row (as inner contrast increases). Instead, gamma was strongest for the contrast matched stimuli along the diagonal compared to stimuli away from the diagonal (Figure 5A). This is also clear in Figure 5B, where each plot represents the change in power for a fixed inner contrast and varying levels of outer contrast, and in the comparison of the mean level of normalized gamma for all stimulus conditions, averaged across all center sites across sessions (Figure 6A). To quantify this effect, we compared the gamma power for each contrast matched stimulus and its neighboring mismatched stimuli, obtained by either decreasing or increasing the contrast of either the inner or outer grating only (Figure 6A, bar plots). Gamma rhythm was negligible for contrast matched stimuli at low contrasts (below 25% in our stimulus set) and therefore power in this band simply followed the change in overall stimulus drive due to discontinuities at low contrasts. However, at higher contrasts (50% and above), any change in contrast of either the center or surround (decrease or increase) from the matched case caused a significant reduction in gamma power. This effect was especially stark when the contrast matched stimulus at 50% or 75% (in Monkey 2) was changed.

**Figure 5:**
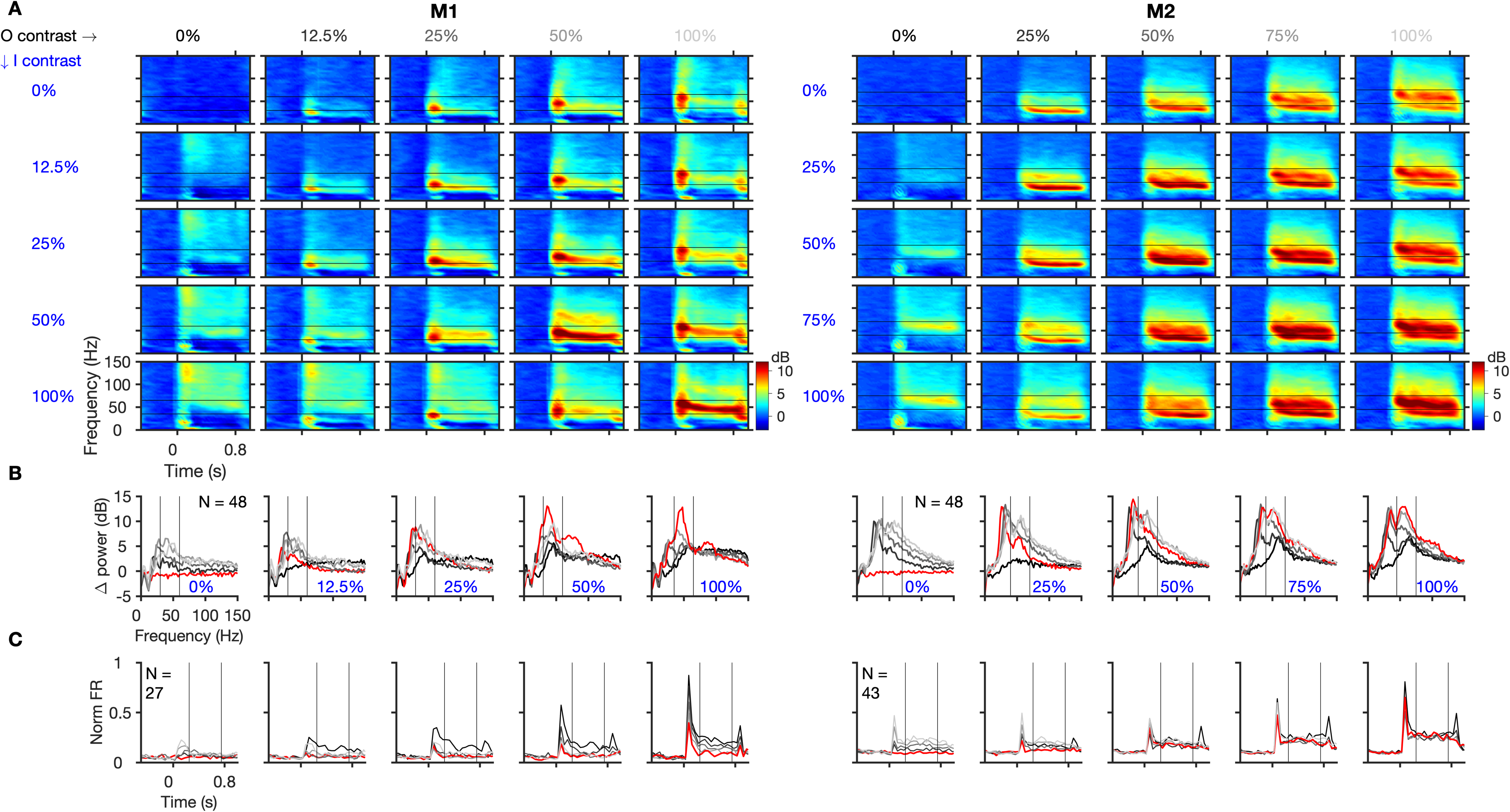
Gamma oscillations are reduced by contrast discontinuity. A) Trial-averaged TF difference spectra for the population, induced by stimuli with contrast discontinuity (inner and outer contrasts are indicated on the left and top respectively) and B) the corresponding mean change in power from baseline to stimulus period. C) Mean normalized firing rate averaged across selected spiking center electrodes and sessions. Same format as Figure 3A-C. In B-C, every column shows graphs for varying outer contrast (increasing with brightness of grey), at a fixed inner contrast (mentioned in (B), and the colored red line represents the stimulus with the corresponding matched inner-outer contrast).

**Figure 6:**
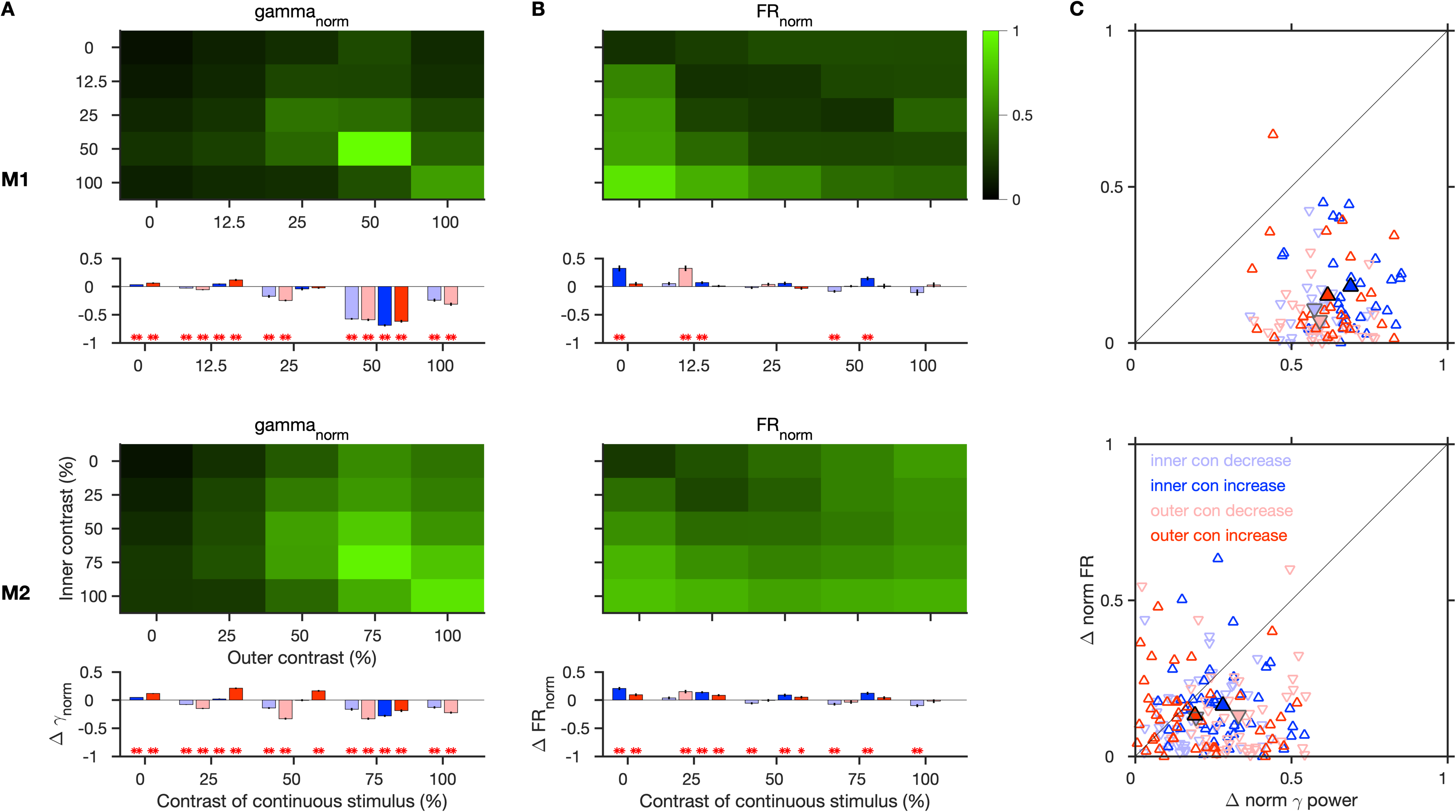
Effect of contrast discontinuity on gamma and firing responses. A) Normalized gamma power and B) Normalized firing rate during the stimulus period for different outer and inner contrast conditions, for the same electrodes as in Figure 5C, for Monkey 1 (top) and 2 (bottom). The corresponding effect of change in contrast from a continuous stimulus is shown below each. The bar graphs denote the change for a step change (decrease or increase) in contrast occurring in either the inner or the outer grating, from a continuous stimulus whose contrast is indicated on the x-axis. Asterisks indicate its statistical significance (** for p<0.01, * for p<0.05, WSR test). C) Magnitude of change in normalized gamma power versus normalized firing rates for the four smallest discontinuities from 50% contrast and 75% contrast continuous grating in Monkey 1 and 2 respectively. Larger and bordered data points show the mean across sites.

The spiking activity, on the other hand, showed a more direct dependence on the contrast, generally increasing with an increase in inner contrast or reduction in outer contrast (Figure 5C, 6B), and decreasing with a decrease in inner contrast. With an increase in outer contrast, the firing rates occasionally increased, especially at low-level center contrasts (0%, 25%) in Monkey 2, although this increase was smaller than the increase due to center contrast (Figure 5C, 6B). Such an effect of surround contrast is not unexpected, since at low contrasts surround has been shown to be facilitatory (Ichida et al., 2007; Polat et al., 1998). At mid and high contrast discontinuities, firing rates mostly remained unchanged.

We compared the effect of contrast discontinuity on gamma and firing rates at each site by plotting the magnitude of change in their normalized values from the continuous case with strongest gamma rhythm (50% in Monkey 1 and 75% in Monkey 2) to the four immediate discontinuity cases i.e. when either the inner or outer contrast changed by one step (Figure 6C). In both the monkeys, the change in gamma was significantly greater than for firing rates in almost all cases (p = 0.56 × 10^−5^, 0.56 × 10^−5^, 0.56 × 10^−5^ and 0.79 × 10^−5^ in Monkey 1 for inner contrast decrease and increase, and outer contrast decrease and increase respectively; p = 0.18 × 10^−2^, 0.11 × 10^−3^, 0.14 × 10^−5^ and 0.78 × 10^−1^ in Monkey, WSR test). Thus, gamma was more sensitive to contrast discontinuities than firing rates.

### A resonant model of gamma oscillations

To understand the resonant properties of gamma oscillations, we used a previously well-studied lumped neuronal model, consisting of an excitatory (E) and an inhibitory (I) neuronal population that can generate oscillations ((Jadi and Sejnowski, 2014), equation 1, refer ‘Model details’ in Methods). To model discontinuities in this framework, we included additional lateral recurrent (LR) inputs (equation 2-3), which were further dependent on the overall excitatory and inhibitory activity of the population (Figure 7). This effectively changed the weights of the network model (see Methods for details). In this framework, loss of recurrent inputs due to a discontinuity could be modeled as a reduction in some of the weights.

**Figure 7:**
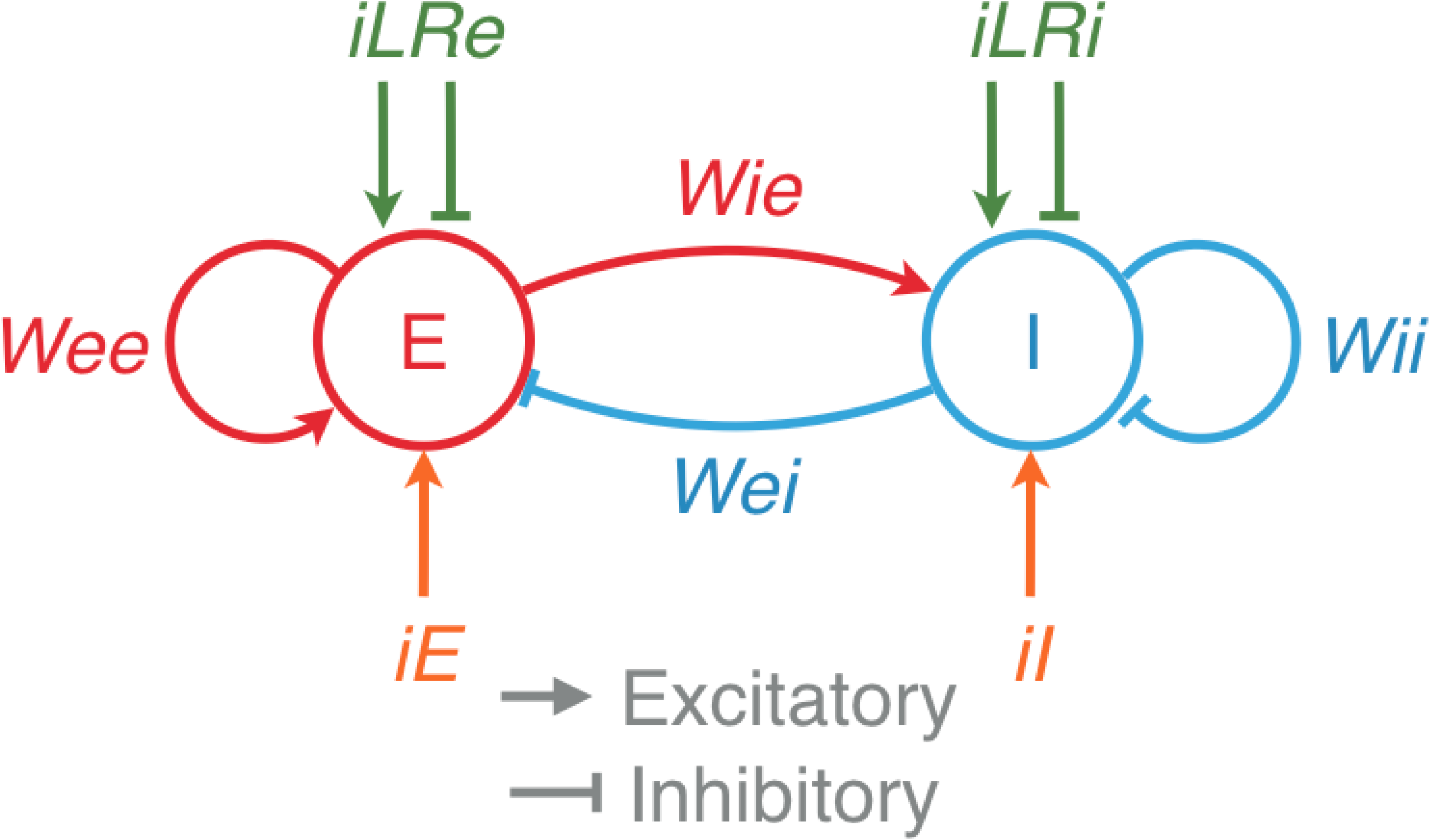
A resonant model of gamma oscillations. A schematic of the model with an excitatory population (E) and an inhibitory population (I). *iE* and *iI* are the external inputs, and *iLRe* and *iLRi* are inputs from other nearby local recurrent networks. *Wee*, *Wei*, *Wie* and *Wii* are the gains of the corresponding connections between the populations.

Figure 8A shows the mean firing rates of the E population (first column), I population (second), peak gamma power (third) and peak gamma frequency (fourth), as a function of the external inputs given to the E and I populations (denoted by *iE* and *iI*). Here, increasing the size of the stimulus increases *iI* (vertical progression in these plots), while increasing the contrast of the stimulus increases both *iE* and *iI* (diagonal progression in the plots). Jadi and Sejnowski showed that in a regime in which the responses of I population is strongly superlinear while E population is sublinear (as indicated by white lines; for details see (Jadi and Sejnowski, 2014)), increasing stimulus contrast increases gamma peak frequency and increasing stimulus size decreases gamma peak frequency and increases its magnitude, as observed in real data (Gieselmann and Thiele, 2008; Henrie and Shapley, 2005; Jia et al., 2013; Ray and Maunsell, 2011, 2010).

**Figure 8:**
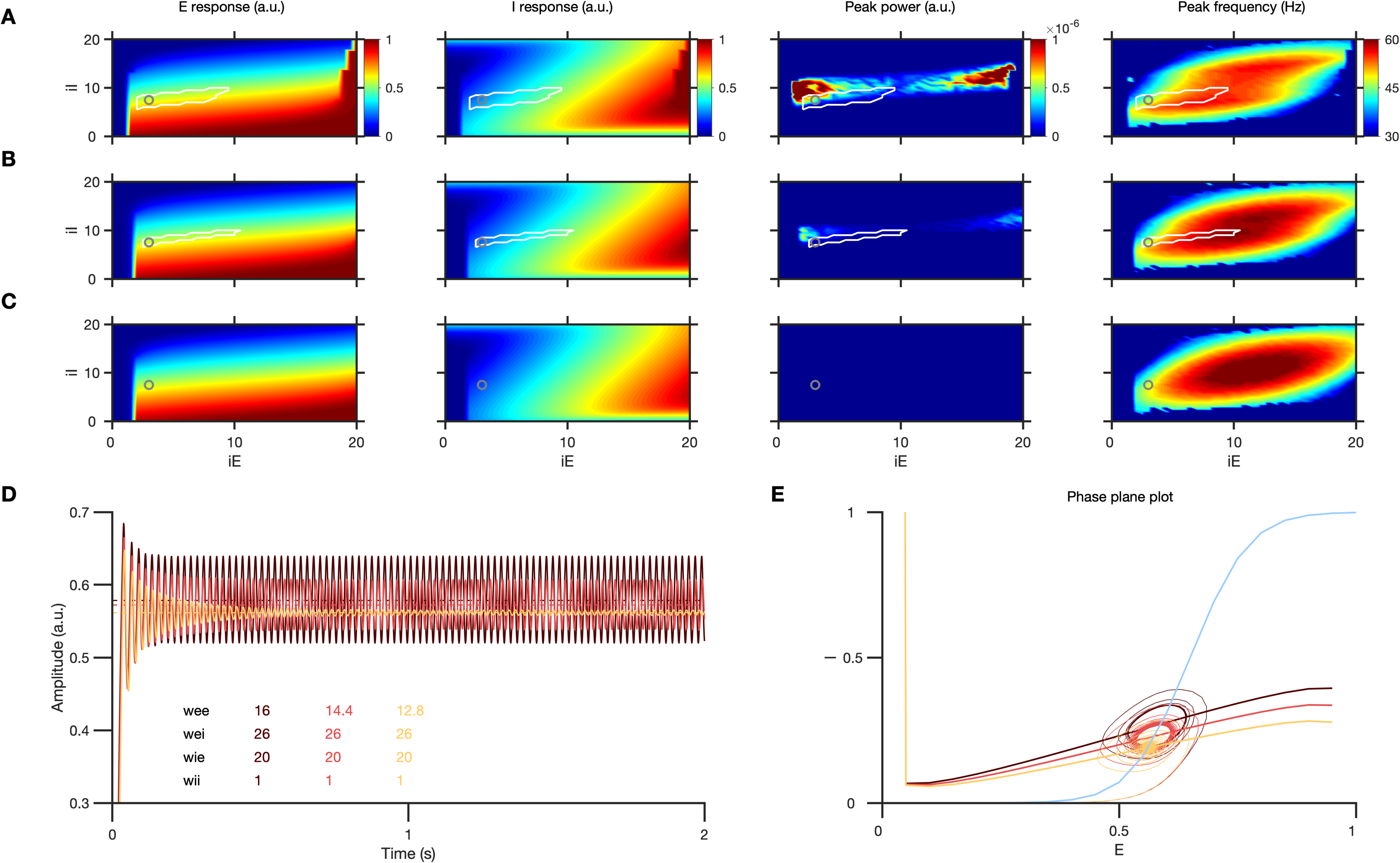
Effect of stimulus discontinuity in the model. A) Left to right: E and I response, peak power in the gamma range and the corresponding peak frequency, for different values if *iE* and *iI* for the default model weights. B) and C) are the corresponding results for modified networks with only *wee* reduced by 10% and 20% respectively, as excitatory LR inputs to E decrease due to discontinuity (weights indicated in D)). White lines indicate the sublinear E and superlinear I response regime where increase in *iI* causes gamma amplitude to increase and frequency to decrease (this regime is absent in case 3). Oscillations weaken drastically in the network from case 1 to 2 and 3, even when the mean E and I responses do not change as much. D) E response in the above 3 cases for *iE* = 3, *iI* = 7.5 (marked by the gray circle). E) Phase-plane analysis showing the effect of varying *wee* on the E and I nullclines. Only the E nullcline shifts in these three cases, while I nullcline (light blue) remains the same.

Here we show that small perturbations in the model operating within this regime, which only lead to small changes in the steady-state firing rates of the E and I populations, can severely attenuate gamma oscillations. We modeled the discontinuity as a drop in the excitatory synaptic weights for E (*wee*) by 10% (Figure 8B) and 20% (Figure 8C). Even a 10% drop in *wee* caused oscillations to weaken considerably (8B, 8D), and they were almost abolished with a 20% reduction (8C-D). Importantly, the mean E or I response remained almost unchanged. To study the properties of this dynamical system, we plotted the excitatory and inhibitory nullclines and found that reduction in *wee* led to a minor flattening of the excitatory nullcline (Figure 8E). While this led to only a minor change in the fixed point (mean E and I responses), the oscillatory power (limit cycle) reduced drastically. Jadi and Sejnowski (2014) had shown that by varying the external inputs (which cause a translation of the E and I nullclines), the dynamical system can transition from a stable operating regime to an oscillatory one through a transition known as a supercritical Andronov-Hopf bifurcation, in which both the power and frequency of the oscillations critically depend on the weights and time constants (exact dependence is derived in Jadi and Sejnowski, 2014 and is therefore not replicated here). Such a finely balanced system near the bifurcation point could be brought back to a stable regime by a slight reduction of the excitatory weights, leading to the loss of oscillations. Modifications in other weights (such as *wii*, *wei* or *wie*), which caused similar changes in the slopes of the E and I nullclines, also led to changes in the oscillatory properties and the expanse of the valid regime where the model behaved as expected. Reduction in weights mostly led to shrinking of this valid regime, and concomitant changes in weights produced even stronger effects. Overall, these results demonstrate a resonance like behavior of the oscillations, similar to the experimental observations, whereby small perturbations (here modelled by a small change in *wee*) that do not change the steady state firing rates by much can nonetheless cause huge changes in the magnitude of the oscillations.

## Discussion

Strong gamma oscillations in V1 LFP are known to be induced by optimized grating stimuli. When the center and surround of V1 RF were stimulated by gratings of different orientations, phases or contrasts, or when there was a grey annular cut between these regions, strength of gamma oscillations was severely reduced. Importantly, gamma oscillations were more sensitive to these discontinuities compared to the spiking activity. Furthermore, this sensitivity of gamma to the discontinuity was dependent on the location of discontinuity with respect to the center-surround structure of the V1 RF. We demonstrate that an E-I neuronal network model can exhibit such a behavior when there are small changes in overall weights in the network due to such discontinuities.

### Gamma and visual irregularities

Gamma oscillations induced by large gratings have been shown to be stronger and coherent over larger spatial extents than those induced by small gratings, but are adversely affected by addition of noise (Jia et al., 2013, 2011; Zhou et al., 2008), or superimposition of cross orientation gratings to form plaids (Lima et al., 2010). While addition of noise or an orthogonal orientation component modifies both the center and surround properties, our manipulations mostly change the stimulus outside the classical RF, bringing out the strong influence of this extra-classical modulation on gamma oscillations. This modulation can be both inhibitory and excitatory, and is mostly mediated through inputs from lateral intracortical connections, feedback from higher areas, and feedforward inputs (Angelucci et al., 2017, 2002; Nassi et al., 2013; Nurminen et al., 2018; Ozeki et al., 2004). In a similar context, a study in mouse V1 (Veit et al., 2017) showed that orthogonal orientation in the extra-classical surround reduces the power of a ‘context-dependent gamma’ rhythm.

Across all the discontinuity types an additional contrast component introduced by the sharp stimulus transition across the discontinuity edge may contribute to the excitation of heterogeneous groups of neurons preferring other orientations and spatial frequency in V1, thereby adding to the E-I imbalance. In this context, the orientation and phase discontinuities have been shown previously to cause an effect called brightness induction or enhancement (Biederlack et al., 2006; Huang et al., 2002; Sillito and Jones, 1996), under which the center appears perceptually brighter due to the orientation contrast or phase contrast. A previous study (Biederlack et al., 2006) reported that the firing rates for the units responding to the center grating increased only with orientation discontinuity but not with phase discontinuity. We found that firing rates increased for both the stimulus variations, as well as for the annular cut case, which has been shown to not cause any brightness enhancement, although the change was not as pronounced as the change in gamma power, especially for the smallest discontinuities we used. In general, however, it has been shown that gamma and spiking activity may not necessarily show a definite relationship across stimulus variations (Jia et al., 2013).

The radial contrast component and the perceptual contrast shift effects as discussed above are studied more directly in the contrast discontinuity experiment. The nature of modulatory inputs from the extra-classical surround region is heavily dependent on the stimulus contrast (Angelucci et al., 2017; Ichida et al., 2007; Polat et al., 1998; Schwabe et al., 2010). The surround region can be suppressive at high contrast but can provide net excitation at a lower contrast. In general, it has been observed that the size of the excitatory RFs in V1 expand slightly at lower contrasts in comparison to the higher contrasts (Cavanaugh et al., 2002; Sceniak et al., 1999). These contrast dependent effects may play a role in modulating the E-I signals in different ways at different contrasts in our stimulus set, leading to slight differences in the effect of discontinuities at different contrasts (Figure 6). The effect of discontinuities is qualitatively similar even when they progressively remove areas of spatial contrast, as in annular discontinuities, modifying the overall stimulated cortical volume itself. While our results pertain to grating stimuli, gamma oscillations have been observed to be reduced by a mismatched surround or annulus for uniform surfaces as well (Peter et al., 2019). These findings suggest that any visual structural modification that changes the critical level and nature of E-I drive in the network can potentially affect gamma.

### Animal specific differences

While all the major results shown here were consistent across monkeys, the magnitude of these effects were in general smaller in Monkey 2 as compared to Monkey 1. One of the factors in this could be that the RFs in Monkey 1 were at a larger visual eccentricity than in Monkey 2. RFs closer to the fovea are known to be smaller, as seen in Monkey 2 here, and the properties of RFs in terms of overall connectivity, gains and center-surround spread and interactions may depend on visual eccentricity. Another notable difference is the presence of a prominent slow gamma rhythm in Monkey 2. The effects of discontinuity on this rhythm were qualitatively similar to fast gamma, although the drop was smaller. In particular, for annular discontinuities at varying inner radii, slow gamma reduced strongly even at farther cuts, which is expected if a spatially more widespread network is involved in its generation (Murty et al., 2018). Moreover, the effect of smallest contrast discontinuity was most drastic for slow gamma in the mid-contrast range, whereas for fast gamma it was in the high-contrast range (Figure 5). This effect incidentally follows the contrast tuning of these two gamma rhythms to large continuous gratings (Murty et al., 2018).

### Implications for natural image processing

Selective gamma synchronization of neuronal ensembles responding to an optimized continuous stimulus may provide stronger or more potent signal for intra and inter-areal communication in the gamma range of frequencies (Buzsáki and Schomburg, 2015; Fries, 2005). However, some other studies have shown gamma to be dependent more on the visual context or the spatial properties of the stimulus with respect to the RF and not necessarily on a unified percept (Dong et al., 2008; Lima et al., 2010; Palanca and DeAngelis, 2005; Thiele and Stoner, 2003). Given the high sensitivity of gamma to small stimulus discontinuities that we show, most natural stimuli might not be able to generate strong gamma oscillations in V1 LFP, as reported by many studies (Bartoli et al., 2019; Hermes et al., 2019, 2014; Kanth and Ray, 2020; Kayser et al., 2003). Gamma is also strongly generated by uniform color surfaces (Shirhatti and Ray, 2018), and therefore some colored natural stimuli may generate strong gamma when favorable uniform color patches fall on the RF (Bartoli et al., 2019; Brunet et al., 2013; Kanth and Ray, 2020). However, mismatches in color surfaces in the form of a differently colored blob or annulus can also reduce gamma power (Peter et al., 2019), suggesting that the effect of discontinuities is qualitatively consistent across both achromatic and chromatic pathways, and that most natural stimuli with intricate structural complexities would not generate strong gamma oscillations.

Studies have shown that the responses in V1 for naturalistic stimuli can be explained by the degree of statistical dependence between the stimulus structures falling on the center and surround regions of the V1 RF (Coen-Cagli et al., 2012; Schwartz and Simoncelli, 2001; Simoncelli and Olshausen, 2001). Considering the high sensitivity of gamma to this visual structure, it will be interesting to determine how gamma depends on natural scene statistics. Recently it has been shown that information about images is high in the LFP gamma band (Kanth and Ray, 2020), and orientation variability in images can be used to predict gamma responses (Hermes et al., 2019). It may be argued that gamma in V1 signifies the stimuli more optimized to V1 response properties and as an extension of this, gamma in higher areas might code for larger surfaces or whole objects more relevant to the response properties of those neurons. However, gamma strength may decrease with increasing eccentricity and RF size leading to less prominent oscillations in higher areas (Vinck and Bosman, 2016).

### Gamma mechanisms and neural models

Neuronal models with recurrent connections between excitatory and inhibitory populations have been used to model and study oscillations in cortical and sub-cortical networks (Jadi and Sejnowski, 2014; Jia et al., 2013; Kang et al., 2010; Kraynyukova and Tchumatchenko, 2018; Roxin and Compte, 2016; Tsodyks et al., 1997; Wilson and Cowan, 1972). Mechanisms of gamma may be understood better by studying how discontinuities affect the overall network dynamics in such models. Cortical neurons are known to have supralinear input-output response functions such that the net synaptic throughput effectively changes with input strength (Ahmadian et al., 2013; Priebe and Ferster, 2008; Rubin et al., 2015). As the network gets engaged more strongly, as in the case of large gratings, normalization signals dominate and inhibition stabilizes the network activity (Heeger, 1992; Miller, 2016; Ozeki et al., 2009). The tuned nature of these signals can modulate network interactions depending on stimulus properties (Angelucci et al., 2017; Ni et al., 2012; Shushruth et al., 2012). Therefore, discontinuities can modulate the overall network drive, which can effectively modify synaptic gains and the operating regime of the network. Incorporation of the detailed RF structure into the model may further unravel mechanistic properties of these oscillations.

In summary, gamma oscillations in V1 LFP exhibit resonance like behavior which could signify critical E-I balance in the neuronal network. Further investigations with finer variations of stimulus properties across this specialized RF structure may lead to better insights into the dynamics and mechanisms of such oscillations. Studying such oscillations in conjunction with spikes and their sensitivity to the spatiotemporal sensory context may help us understand general principles of cortical sensory processing.

## MATERIALS AND METHODS

### Animal preparation and training

All experiments were performed as per the guidelines approved by Institutional Animal Ethics Committee (IAEC) of the Indian Institute of Science and the Committee for the Purpose of Control and Supervision of Experiments on Animals (CPCSEA). For this study two adult female monkeys (Macaca radiata; 13 years, ∼3.3 Kg; 17 years, ∼4 Kg) were used. Animal preparation and training details are the same as described in earlier studies (Murty et al., 2018; Shirhatti and Ray, 2018). Each monkey was surgically implanted with a titanium headpost over the anterior/frontal region of the skull under general anesthesia. Following recovery, the monkey was trained on a passive visual fixation task, and once the monkey reached a satisfactorily level of training, another surgery was performed under general anesthesia to insert a microelectrode array (Utah array, 96 active platinum microelectrodes, 1 mm length, 400 µm inter-electrode distance, Blackrock Microsystems) in the primary visual cortex (right hemisphere, centered at ∼10-15 mm rostral from the occipital ridge and ∼10-15 mm lateral from the midline; the location varied slightly across the two monkeys). The receptive fields of the recorded neurons were located at an eccentricity of between ∼3° to ∼4.5° in Monkey 1 and between ∼1.4° to ∼1.8° in Monkey 2, in the lower left quadrant of the visual space with respect to fixation. Considering the dimensions of the microelectrodes, the recordings are most likely to be around cortical layer 2/3.

### Experimental setup and behavior

Details of experimental setup, behavior and data recording are as described in previous studies (Murty et al., 2018; Shirhatti and Ray, 2018). Each monkey viewed a monitor (BenQ XL2411, LCD, 1280 × 720 resolution, 100 Hz refresh rate, gamma corrected and calibrated to a mean luminance of 60 cd/m^2^ on the monitor surface using i1Display Pro, x-rite PANTONE) placed ∼50 cm from its eyes, with its head fixed by the headpost in a custom designed monkey-chair. A Faraday enclosure (constructed using thin copper sheets, wood and sound isolating material), with a dedicated grounding separate from the mains supply ground, was used to house the monkey chair and the display and recording setup during experiments.

The monkeys performed a passive visual fixation task, in which they had to maintain visual fixation at a small dot (0.05° or 0.10° radius) at the center of the screen for the duration of a trial, which could be either 3.3 or 4.8 s. Each trial began when the monkey fixated, and following an initial blank grey screen of 1000 ms, 2 to 3 stimuli were shown for 800 ms each, with an inter-stimulus interval of 700 ms. The monkey was rewarded with juice for successfully holding its fixation without blinking, within 2° of the fixation spot, which stayed on throughout this period. Although the fixation window was kept slightly large, mainly to adjust for occasional small shifts in the head position due to slight movements of the chair and related apparatus, the monkeys actually maintained fixation within a much smaller window during the task. The standard deviation of eye position during a trial across sessions was small on an average for both monkeys (< 0.18° and < 0.16° along the horizontal and vertical axes respectively for Monkey 1; < 0.28° and < 0.29° for Monkey 2).

### Data recording

A 128-channel Cerebus Neural Signal Processor (Blackrock Microsystems) was used to record raw signals on 96 channels. To obtain the LFPs, these raw signals were filtered online between 0.3 Hz (Butterworth filter, 1^st^ order, analog) and 500 Hz (Butterworth filter, 4^th^ order, digital), sampled at 2 KHz and digitized at 16-bit resolution. To extract multiunit spikes, the raw signals were filtered online separately between 250 Hz (Butterworth filter, 4^th^ order, digital) and 7.5 KHz (Butterworth filter, 3^rd^ order, analog) and the filtered signal was subjected to a threshold (amplitude threshold of ∼5 standard deviations of the signal). No further offline filtering of the LFP signals or offline spike sorting was done.

An ETL-200 Primate Eye Tracking System (ISCAN Incorporated) was used to record eye position data in terms of horizontal and vertical co-ordinates/position and pupil diameter, at a sampling rate of 200 Hz during the task. A custom software running on MAC OS monitored the eye signals, and controlled the progression of task and trials, stimulus generation and pseudorandom stimulus presentation.

### Electrode selection and data analysis

For each monkey, a receptive field mapping experiment was run regularly across multiple sessions across days to verify the stability of RFs and assess the suitability of electrodes for data analyses. In this experiment, small sinusoidal gratings (radius of 0.3° and 0.2° for Monkey 1 and 2 respectively; static, full contrast, spatial frequency of 4 cycles per degree (cpd), at four orientations of 0°, 45°, 90° and 135° in both monkeys) were flashed for 200 ms at equally spaced (9 × 9) locations within a rectangular grid on the visual space that approximately covered the aggregate RF of the entire microelectrode array. A two-dimensional Gaussian fitted to the spatial profile of the maximum drop in the LFP (during 40-100ms after stimulus onset for both monkeys; baseline corrected) induced by stimuli presented at the various locations yielded the RF estimate for each electrode (Dubey and Ray, 2016). Electrodes with consistent stimulus induced changes in LFP and reliable estimates of RF size across sessions were chosen after subjecting their mean distribution across sessions to an arbitrary threshold based on inspection. Since we primarily analyzed and characterized LFP responses, an LFP based measure was used, although estimates based on spiking activity were similar (Dubey and Ray, 2019, 2016). Electrodes which showed noisy or inconsistent signals; or a high degree of crosstalk across sessions; or impedances outside the range 250 KΩ to 2500 KΩ, were discarded from analyses. This procedure yielded an overall 65 and 39 usable electrodes for Monkey 1 and 2 respectively, which were considered for all further analyses. Out of these, the electrodes that showed high impedance during certain sessions were not considered for analysis for that session. The resultant set of electrodes were considered while choosing the center sites/electrodes (explained ahead) during any session. The final number of electrodes used for analysis of data from different sessions are stated in the figures or in the related text descriptions.

For each session the sites whose RFs were sufficiently close to the stimulus center (within a range of 0.2° and 0.15° for monkeys 1 and 2 respectively; a smaller range was chosen for Monkey 2 since the RFs were less eccentric and smaller than in Monkey 1) were selected as the ‘center’ sites for analyses. For the annular cut discontinuity experiment with different cut locations, this range was 0.15° and 0.1° for monkeys 1 and 2 respectively, corresponding to the nearest cut location. All analyses were done for such sites and the mean was taken across sites across sessions.

Every experiment was repeated over several sessions, each yielding data from a few sites whose RFs were close to the center of the stimulus. After rejection of electrical artifacts or noisy data (average rejection rate: 1.57 % in Monkey 1, 2.07 % in Monkey 2; procedure summarized in (Murty et al., 2018)), the number of sessions, number of center electrodes, and the mean number of repeats across sessions, for the two monkeys respectively, were as follows - Annular cut discontinuity experiment (Figure 1): 13 sessions, 60 electrodes (out of which 44 were unique, since some electrodes were repeated across sessions), 11.02 repeats for Monkey 1 and 8 sessions, 53 (32 unique) electrodes, 15.16 repeats for Monkey 2; Annular cut discontinuity experiment with different cut locations (Figure 2, Supplementary Figure 1): 13 sessions, 33 (25 unique) electrodes, 11.93 repeats for Monkey 1 and 7 sessions, 21 (17 unique) electrodes, 15.23 repeats for Monkey 2; Orientation discontinuity experiment (Figure 3): 11 sessions, 57 electrodes (43 unique) and 12.32 repeats for Monkey 1 and 9 sessions, 74 electrodes (32 unique) with 14.55 repeats for Monkey 2; Phase discontinuity experiment (Figure 4): 12 sessions, 62 (44 unique) electrodes, 12.42 repeats for Monkey 1 and 13 sessions, 124 (37 unique) electrodes, 16.09 repeats for Monkey 2; Contrast discontinuity experiment (Figure 5-6): 9 sessions, 48 (42 unique) electrodes, 11.95 repeats for Monkey 1 and 6 sessions, 48 (27 unique) electrodes, 14.72 repeats for Monkey 2. All these data analyses were performed using custom written codes in MATLAB, MathWorks.

#### Spectral Analyses

Spectral analyses were performed using the Multitaper method, implemented using functions in the ‘Chronux toolbox’(Mitra and Bokil, 2008) (http://chronux.org/, developed for MATLAB). To obtain the time-frequency difference spectrum, the power spectrum was first calculated using a single taper with a sliding window of 0.25s yielding a 4 Hz frequency resolution. The logarithm of the mean power spectrum across repeats was computed for each electrode. Then the mean power at each frequency in the mean spectrum during the spontaneous period (0.5 to 0 s before stimulus onset) was calculated and subtracted from the entire spectrum, followed by multiplication by 10 to get the difference spectrum in decibel.

Power spectral density (PSD) were computed using a single taper between 0.25-0.75 seconds after stimulus onset and compared to the PSD during spontaneous period (0.5 to 0 seconds). The change in power was calculated by subtracting the logarithm of power during stimulus period and spontaneous activity and multiplying by 10 to yield units in decibels. In Figure 8, PSD was similarly calculated using a single taper for the signal between 1s to 2s.

Power in any frequency band is calculated as the sum of PSD at all frequencies in that band. Normalized gamma power was calculated for each electrode by first calculating the mean gamma power for each condition and then dividing these by their maximum value across conditions for that electrode. The change due to the smallest discontinuity was computed by subtraction of the normalized gamma power for the continuous stimulus from that for the smallest discontinuity stimulus. In the case of orientation and phase discontinuities an average of values for the smallest discontinuities on either side was considered.

Large gratings that substantially extend into the surround region of the RF can induce two distinct gamma rhythms (slow and fast), whose strength and peak frequency can vary with the grating orientation (Murty et al., 2018). Since the stimuli in this study were optimized to generate strong fast gamma rhythms, analyses and results are focused on this band. Therefore, across all figures and results ‘gamma’ refers to fast gamma unless stated otherwise. For all static grating stimuli at 100 % contrast (Figure 1 to Figure 4), fast gamma band was chosen as per inspection of PSD as: [35 65] Hz in Monkey 1, and [45 75] Hz in Monkey 2. The same bands were chosen in the contrast discontinuity experiment, for stimuli in which at least one of the inner or the outer grating was at 100 % contrast. For others, in which both inner and outer grating were at a contrast lower than 100 %, the bands were shifted lower by 5 Hz, since gamma rhythm is known to shift lower in frequency with decreasing contrast, and as also observed in the corresponding PSD.

#### Analyses of spiking units

The center units which showed clear spikes and had an average firing rate of at least 1 spike/second for at least one of the stimulus conditions were chosen for analyses of firing responses. The firing rates for each unit were normalized by dividing by the maximum firing rate for that unit across time and conditions to obtain the mean normalized firing rate for each condition. For comparison of normalized gamma power and normalized firing rate these same spiking units were considered. The mean normalized value of gamma power and firing rate for each condition were calculated by averaging these quantities across the chosen units.

### Model details

The model consists of two interconnected neuronal populations representing a local recurrent network in an orientation column of V1, with external inputs to the network arising from stimulation and lateral or feedback signals. If E and I are considered as the response variables representing the mean response of the two neuronal populations, then the equations governing the dynamics of E and I are given by,

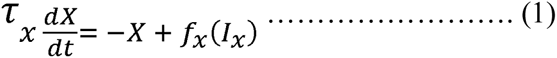

Where *x* can be E or I. *τ_x_* is the corresponding time constant of the variable build up, and *f_x_* is the function transforming the total synaptic input, *I_x_*, to the resultant spiking response.

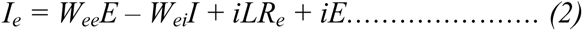

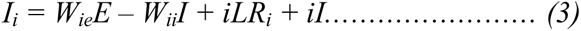

*f_x_* is considered to be a sigmoid function as described in (Jadi and Sejnowski, 2014).

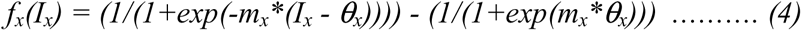

The external inputs arising from the center (inner) and surround (outer) stimuli are *iE* and *iI* for E and I respectively. These equations are similar to the model used by Jadi and Sejnowski (Jadi and Sejnowski, 2014), except there were no separate lateral inputs (*iLR_e_* and *iLR_i_*) in their model. We added *iLR_e_* and *iLR_i_* as the effective intracortical inputs from similar nearby local recurrent input networks through interconnections in the orientation hypercolumn (Angelucci et al., 2017; Shushruth et al., 2012). *W_xy_* implies the synaptic gain from input *y* to the receptor *x*. Considering that all nearby local recurrent networks differ only in their orientation tuning, *iLR_x_* is effectively a weighted function of the E and I activity, for the sake of this simplified single column model design. Therefore equations 2 and 3 can be rewritten as follows.

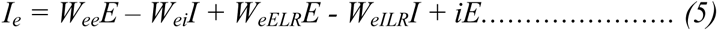

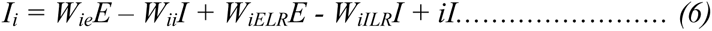

*W_xELR_* and *W_xILR_* denote the gain function for the excitatory and inhibitory inputs respectively from the other local recurrent networks in the hypercolumn. Note that in a multi-column architecture these inputs (*iLR_x_*) can be derived from the actual outputs of the nearby columns. Equations (5) and (6) can be written as follows.

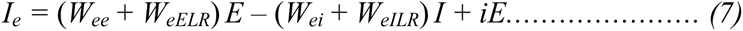

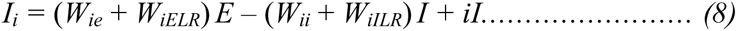

In this model discontinuities can effectively decrease *W_xELR_* and/or *W_xILR_*, and thereby the overall network gains, pushing the network into a different regime of operation.

Equations (7) and (8) can be summarized as follows.

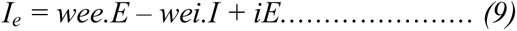

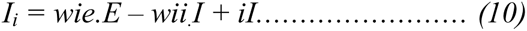

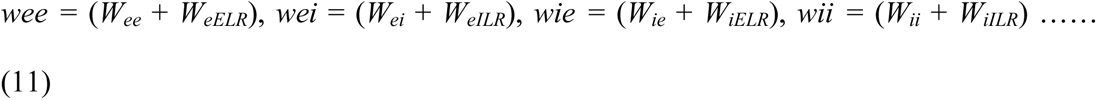

The default values of parameters, adapted from (Jadi and Sejnowski, 2014), are as follows. *wee* = 16, *wei* = 26, *wie* = 20, *wii* = 1, *m_E_* = 1, *m_I_* = 1, *θ_E_* = 5, *θ_I_* = 20, τ_E_ = 20, τ_I_ = 10. These parametric values have been shown to generate gamma oscillations in the network whose properties (peak power and frequency) agree with experimental observations under certain regimes of the network operation, and the network behaves as an inhibition stabilized network (ISN) (Jadi and Sejnowski, 2014; Ozeki et al., 2009). In Figure 8, only *W_eELR_* is considered to be reduced by a discontinuity such that (*W_ee_* + *W_eELR_*), i.e. *wee*, effectively decreases by 10% and 20%, and the network moves drastically to a regime with weak oscillations under new weights. To determine the regime where increase in *iI* caused an increase in the magnitude of oscillation and decrease in its peak frequency, we followed the conditions derived in (Jadi and Sejnowski, 2014) (Appendix B, equation 6, 7, 12, 13 and 14) and the procedure described therein.

## Acknowledgements

This work was supported by Wellcome Trust/DBT India Alliance (IA/S/18/2/504003 – Senior Fellowship to S.R and 500145/Z/09/Z – Intermediate fellowship to SR) and DBT-IISc Partnership Programme.

## Competing financial interests

The authors declare no competing financial interests.

---

**Supplementary Figure 1:**
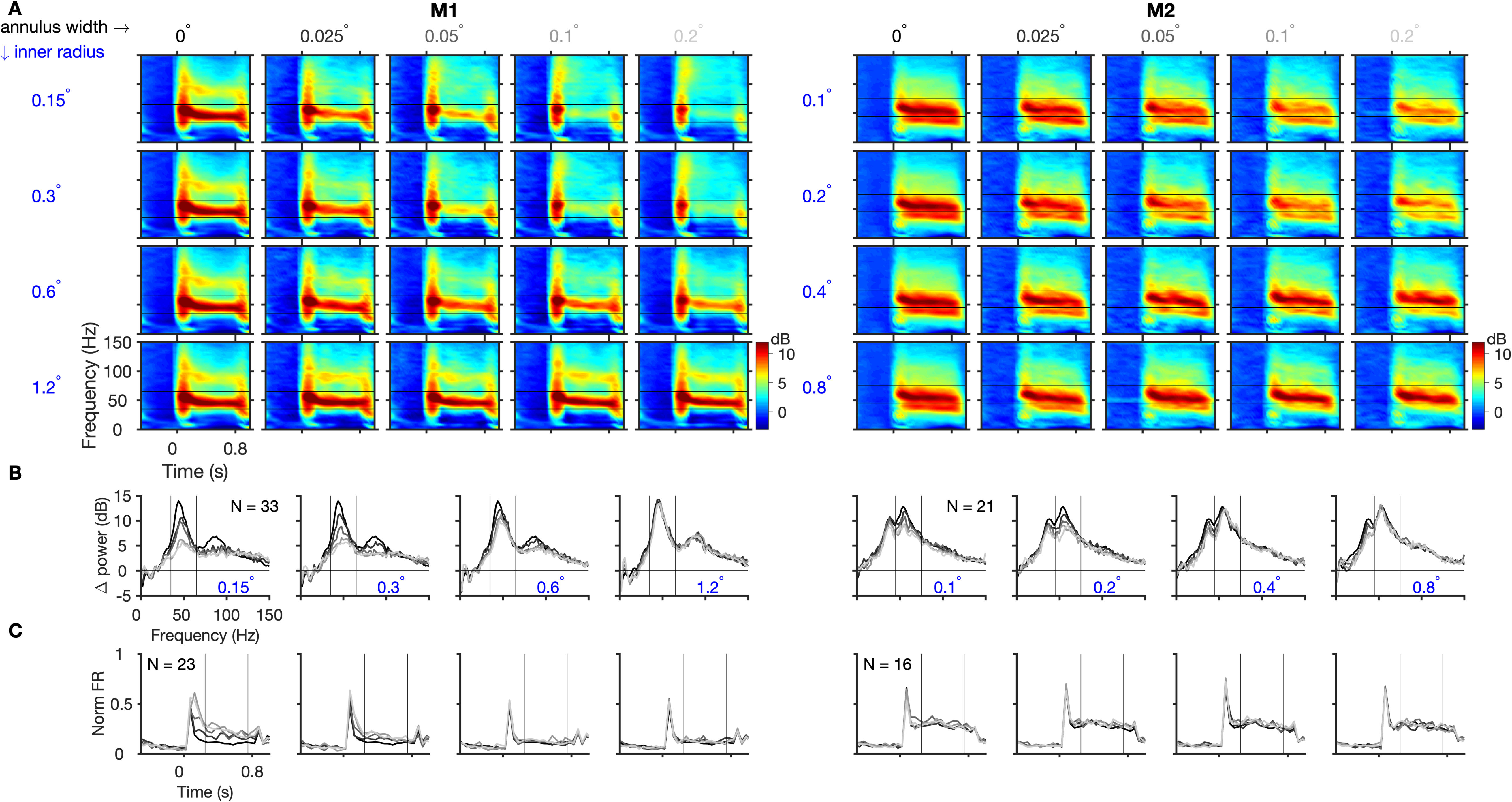
Effect of annular discontinuity at different locations. A) Trial-averaged time-frequency difference spectra for the population in Monkey 1 (left column) and Monkey 2 (right column), induced by stimuli with annular discontinuity at different inner radii (inner radius and annulus width are indicated on the left and top respectively) and B) The corresponding mean change in power from baseline to stimulus period. C) Mean normalized firing rate averaged across selected spiking center electrodes and sessions.

## Notes

### Competing Interest Statement

The authors have declared no competing interest.

